# The Mitochondria-Targeted Peptide HDAP2 Reduces Mitochondrial Loss and Retinal Ganglion Cell Degeneration After Optic Nerve Injury

**DOI:** 10.1101/2023.09.28.560029

**Authors:** Margaret A. MacNeil, Sara Arain, Widnie Mentor, Virginia Garcia-Marin, Alexander Birk

## Abstract

Mitochondrial dysfunction is a critical early driver of retinal ganglion cell (RGC) loss in optic nerve injury. We evaluated whether HDAP2, a mitochondria-targeted aromatic peptide designed to support mitochondrial membrane integrity, could preserve neuronal structure after optic nerve crush (ONC). Systemically administered HDAP2 penetrated the blood–retinal barrier and localized to RGCs and mitochondrial-rich retinal layers. Daily treatment (IP, 3 mg/kg for 14 days) significantly improved RGC survival compared to saline-treated ONC animals. RGC densities increased across central, midperipheral, and peripheral regions, and surviving RGCs exhibited approximately two-fold higher RBPMS expression, suggesting improved cellular health.

Transmission electron microscopy revealed that HDAP2 substantially reduced mitochondrial loss within crushed optic nerve axons. Mitochondrial density in HDAP2-treated nerves reached 70% of uninjured levels (0.129 ± 0.029 vs. 0.185 ± 0.017) and was 5.3-fold higher than untreated ONC nerves (0.025 ± 0.012; Kruskal–Wallis p = 0.039; Cohen’s d = 2.77). Mitochondrial morphology was similar across groups, indicating that HDAP2 prevents mitochondrial loss rather than rescuing damaged organelles. HDAP2-treated nerves also contained a higher density of structurally intact axons, consistent with reduced ultrastructural degeneration following injury.

These findings demonstrate that HDAP2 limits mitochondrial loss and attenuates neuronal degeneration after ONC. Together, the results support HDAP2 as a promising therapeutic candidate for protecting CNS projection neurons by maintaining mitochondrial stability after axonal injury.

## Introduction

Degeneration of retinal ganglion cells (RGCs) and their axons is a hallmark of optic neuropathies, including glaucoma (Jassim et al., 2021; Nickells et al., 2012; Williams et al., 2020) traumatic optic neuropathy (Au & Ma, 2022) and diabetic retinopathy (Ren et al., 2022). The cause of these degenerations is unknown, but likely results from multiple factors, including aging (Hou et al., 2022; Samuel et al., 2011), impaired axonal transport (Crish et al., 2010), and loss of neurotrophic support (Almasieh et al., 2012), but mitochondrial dysfunction plays a particularly significant role (Schrier & Falk, 2011). As mitochondria generate ATP needed for cellular activities, their dysfunction results in excess production of reactive oxygen species (ROS), which initiates a harmful cascade of events that can lead to dendritic atrophy, axonal degeneration, and ultimately neuronal death.

RGCs are among the most metabolically active neurons in the central nervous system and yet have very limited energy reserves when challenged. Cytoplasmic space for excess mitochondria is restricted in RGCs, and mitochondria are strategically positioned within compartments where energy demand is highest (Perge et al., 2009), including at synapses (Faits et al., 2016), in the vicinity of voltage gate sodium channels (Barron et al., 2004) and in unmyelinated axons near the optic nerve head (Bristow et al., 2002; Wilkison et al., 2021). Optic neuropathy-related cellular challenges, such as axonal transport disruption, ischemia/reperfusion, ocular hypertension, and inflammatory signaling, can rapidly perturb mitochondrial homeostasis, leading to mitochondrial membrane depolarization, mitochondrial transport impairment, cristae remodeling, and ROS elevation (Ito & Polo, 2017). These local energetic deficiencies could, in turn, rapidly activate the intrinsic apoptotic pathway of mitochondrial outer-membrane permeabilization, cytochrome c release, and caspase-9/3 activation, which culminates with dendritic atrophy, axonal degeneration and neuronal death (Bossy-Wetzel et al., 1998; Oshitari et al., 2008; Wang et al., 2021). We sought to determine whether a treatment that maintains mitochondrial stability during injury could interrupt this pathological cascade and attenuate RGC loss.

Several new neuroprotective strategies that have focused on enhancing mitochondria function have shown success. One promising approach involves nicotinamide supplementation to maintain NAD⁺ levels, which have been effective in protecting RGCs in glaucoma (Tribble et al., 2021; Williams et al., 2017). Another strategy, transplantation of mitochondria into the eye, also enhances survival of RGCs after injury (Nascimento-Dos-Santos et al., 2020). However, the benefit of this procedure is short-term and introduces procedural risks and costs that limit long-term clinical feasibility.

Mitochondria-targeted peptides have also shown potential in mitigating the effects of mitochondrial dysfunction. Compounds such as SS-31 (Birk et al., 2013; Szeto, 2014), SkQ (Skulachev et al., 2009), and MitoQ (Murphy & Smith, 2007) protect mitochondria by scavenging ROS, thereby limiting the downstream effects of oxidative damage. SS-31 has demonstrated some efficacy in protecting RGCs in a rat glaucoma model (Wu et al., 2019), but methodological limitations of the study, including small sample sizes and insufficient detail regarding retinal sampling locations, make it difficult to assess the robustness of the observed effects. Other mitochondria-targeted peptides have faced additional challenges including poor bioavailability, proteolytic degradation, and toxicity concerns (Constance & Lim, 2012).

HDAP2 was developed to address these limitations by targeting cardiolipin-rich mitochondrial membranes with high affinity. Prior biochemical studies demonstrate that HDAP2 interacts with mitochondrial membranes and enhances their stability under stress (Birk et al., 2025). Unlike peptides that act primarily as ROS scavengers, HDAP2 is designed to support mitochondrial membrane architecture during injury.

While detailed biochemical characterization of HDAP2-cardiolipin interactions has been described elsewhere (Birk et al., 2025), the current study focuses on therapeutic efficacy and safety in the optic nerve crush (ONC) model. We administered 3 mg/kg/day HDAP2 intraperitoneally (IP), well below the toxic range, to test the ability of HDAP2 to prevent RGC loss in vivo after ONC. We quantified RGC survival using both absolute densities and RBPMS:ChAT ratios and used transmission electron microscopy to determine whether HDAP2 reduces mitochondrial loss within injured optic nerve axons.

## Experimental Procedures

### Peptide synthesis and structure

We designed a novel, high-density aromatic peptide (HDAP2; Biotin-dArg-Phe-Phe-dArg-amide) to support mitochondria under conditions of cellular stress. HDAP2 was synthesized commercially (GenScript, Piscataway, NJ, USA) and consists of D-arginines flanking a Phe-Phe dipeptide core, amidated at the C-terminus, and biotinylated at the N-terminus for detection. The peptide binds cardiolipin on the inner mitochondrial membrane and, in cultured cells, supported mitochondrial potential during serum starvation and conditions of elevated oxidative stress (Birk et al., 2025). Peptide structure was verified by HPLC, and mass spectrometry confirmed that purity exceeded 96%. The peptide is patented through the Research Foundation of the City University of New York.

### Animal procedures

We studied C57BL/6 mice (2–6 months, both sexes) that were bred and housed at York College, CUNY. All procedures were approved by the York College IACUC and complied with the ARVO Statement for the Use of Animals in Ophthalmic and Vision Research. Mice were housed under standard laboratory conditions (12-h light/dark cycles) with food and water available ad libitum.

Altogether, 82 animals were used in this study to evaluate HDAP2 toxicity (n=48), uptake of HDAP2 in the retina (n=3), and to assess RGC survival and mitochondrial density following ONC (n=31). In the ONC groups, mice were randomly assigned to receive either HDAP2 (3 mg/kg/day, IP for 14 days, (n=17) or saline vehicle (n=14). All ONC animals underwent unilateral ONC with the contralateral eye used as an uninjured control. An additional 14 animals received either HDAP2 (n=8) or saline vehicle (n=6) without ONC to serve as uninjured controls for establishing baseline RGC densities and RBPMS:ChAT ratios, and to verify that HDAP2 exposure alone does not affect RGC survival in the absence of injury.

### Toxicity evaluation of HDAP2

We assessed the safety of HDAP2 in C57BL/6J mice (2–6 months, both sexes). Animals received daily injections of the peptide at fixed doses for 14 days. Groups with fixed doses were used to determine the LD₅₀ and the maximum tolerated dose (Table 1). Animals were observed twice a day for body weight loss (>10%), behavioral alterations (grooming, exploration), sedation, seizure activity and death. Negative effects were rated on a 0-5 scale in accord with the OECD Test Guideline 425 (OECD, 2022): 0 = no observable effect; 1 = sedation <60 min; 2 = sedation 60-180 min; 3= sedation >180 min; 4 = generalized seizure; 5 = death. Toxicity values were determined using Kaplan–Meier survival and toxicity curves. LD₅₀ values were calculated with nonlinear regression for 14 days of survival. Composite toxicity scores were calculated for each animal by day, normalized to the survival interval, and then averaged by cohort. LD₅₀ values were determined by nonlinear regression using mean toxicity scores for each dose group. All tests were two-tailed and significance was accepted when p < 0.05.

**Table 1.**
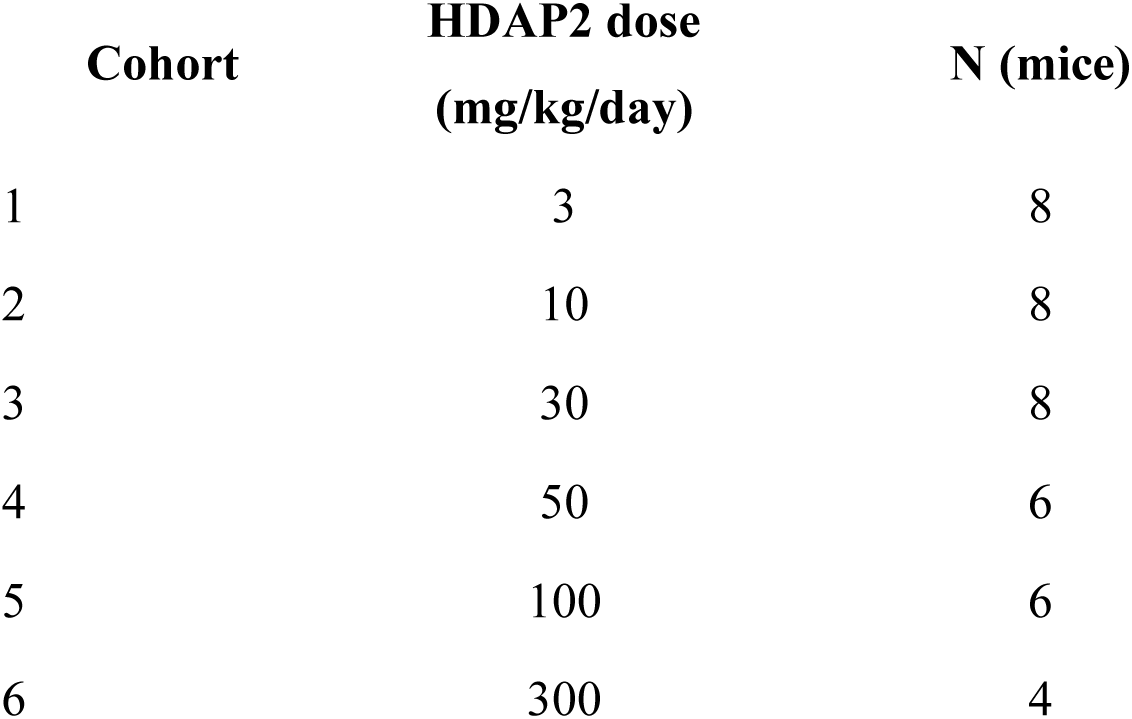
HDAP2 Dosing Schedule for Toxicity Assessment.

| Cohort | HDAP2 dose<br>(mg/kg/day) | N (mice) |
| --- | --- | --- |
| 1 | 3 | 8 |
| 2 | 10 | 8 |
| 3 | 30 | 8 |
| 4 | 50 | 6 |
| 5 | 100 | 6 |
| 6 | 300 | 4 |

### Distribution of HDAP2 in retina

Prior to ONC experiments, we verified that biotinylated HDAP2 was colocalized with RGCs in the retina. Adult mice were injected with 50 mg/kg biotinylated HDAP2 (IP) reconstituted in saline and observed for 2 hours before euthanizing. This higher dose was used during initial toxicity assessment to maximize tissue concentrations of HDAP2 for immunodetection, as HDAP2 exhibits rapid cellular uptake and clearance kinetics. The therapeutic efficacy studies (described below) used a lower dose (3 mg/kg/day, IP), which was adequate to provide neuroprotective effects while still maintaining a wide safety margin (LD50 = 55 mg/kg). Eyes were marked to record orientation, and animals were euthanized with an overdose of ketamine (300 mg/kg) and xylazine (60 mg/kg) administered IP before removing each eye from the orbit with forceps.

Whole eyes were immersed in 4% paraformaldehyde reconstituted in either 0.1 M Tris or phosphate buffer (pH 7.3). After 10 minutes, the cornea, lens, and vitreous body were removed to ensure better penetration of the fixative into the retina, and the eyecups were returned to fixative for 1 hour. Following fixation, retinas in the eyecup were rinsed in buffer, cryoprotected in 0.1 M Tris or phosphate-buffered sucrose (10%, 20%, and 30%) prior to embedding in optimal cutting temperature (OCT) medium (Polyfreeze, Polysciences, Warrington, PA). Sections were cut at 14 μm thickness at −25°C using a cryostat (Leica CM3050S) and mounted onto gelatin coated slides.

HDAP2 uptake was visualized by incubating tissue overnight with Alexa Fluor-conjugated streptavidin (594 nm, 1:500, Jackson ImmunoResearch, West Grove, PA). Tissues were co-stained with antibodies for mitochondrial Cox4, RBPMS (RNA-binding protein with multiple splicing, which labels retinal ganglion cells), and glutamine synthetase (which labels Müller cells), as described below.

### Optic nerve crush procedure

Adult mice (2–6 months) were anesthetized with ketamine (85 mg/kg) and xylazine (20 mg/kg) administered IP. The corneas were numbed with 0.5% proparacaine hydrochloride ophthalmic solution (Falcon Pharmaceuticals, Fort Worth, TX), and animals were placed on their right side under a Nikon dissecting microscope with the head rotated so the snout was facing 2 o’clock, which allowed optimal visualization and access to the dorsomedial conjunctiva of the left eye.

The dorsolateral conjunctiva was cut with microdissection spring scissors, the intraocular muscles were separated, and the eyeball was slightly retracted to expose the optic nerve. Fine, self-closing forceps (RS-5027, Roboz, Gathersburg, MD) were used to clamp the optic nerve 1–2 mm behind the globe for approximately 3 seconds without applying additional tension (Kalesnykas et al., 2012; Tang et al., 2011). After the crush, the eye was repositioned within the orbit, the conjunctiva reflected over the extraocular muscles, and the incision covered with ophthalmic polymyxin B-neomycin-bacitracin ointment (3.5 mg/g, Bausch and Lomb, Bridgewater, NJ). The optic nerves from the contralateral eye were untouched and used as untreated controls.

Following the crush procedure, mice were given a subcutaneous injection of extended-release buprenorphine (0.05 mg/kg, Ethiqua XR) for analgesia and placed on a heating pad until awake and grooming, at which point they were returned to their home cages. Half the animals were given HDAP2 immediately following the procedure and then daily thereafter (3 mg/kg, IP) for 14 days

### Tissue collection and processing

To assess RGC survival and mitochondrial density in the optic nerve, animals were euthanized 14 days post-ONC with an IP overdose of ketamine (300 mg/kg) and xylazine (60 mg/kg). Each eye was removed from the animal with forceps and hemisected. For retinal analysis, whole eyes were fixed in 4% paraformaldehyde for 1 hour as described in the Immunohistochemistry section below. For mitochondrial analysis, optic nerves were transected at the optic nerve head and immersed in 2.5% glutaraldehyde, 2% paraformaldehyde until embedding for electron microscopy.

### Immunohistochemistry

Fixed retinas were labeled with cell markers using conventional immunohistochemical protocols to identify cells and structures associated with HDAP2, and to assess retinal ganglion cell and starburst amacrine cell survival following optic nerve crush.

Retinas and sections were blocked and permeabilized with 4% normal donkey serum in 0.1 M Tris or phosphate buffer with 0.5% Triton-X for 1 hour, then incubated overnight (sections) or for 3 days (wholemounts) at 4°C with selected primary antibodies in 2% normal donkey serum in the same buffer. Tissue was rinsed 3 × 10 minutes with buffer and incubated with secondary antibodies for 1 hour at room temperature (sections) or overnight at 4°C (wholemounts).

Primary antibodies used were: rabbit anti-RBPMS (1:200; PA5-31231, Invitrogen, Carlsbad, CA) to label all retinal ganglion cells; goat anti-choline acetyltransferase (ChAT, 1:100; AB144P EMD Millipore, Burlington, MA) to label starburst amacrine cells; rabbit anti-Cox4 (1:200; PA5-29992, Invitrogen, Carlsbad, CA) to identify mitochondria; and glutamine synthetase (which labels Müller cells, 1:300, Invitrogen, # PA1-46165, Carlsbad, CA). Secondary antibodies used a donkey host and were conjugated with Alexa Fluor 488, 594, or 647 (1:500; Life Technologies Corp., Carlsbad, CA). To identify cell nuclei and delineate retinal layers, tissue was labeled with ToPro3 nuclear stain (1:1000; Invitrogen, Carlsbad, CA) for 15 minutes. After staining, sections and retinas were rinsed in buffer and coverslipped with Vectashield mounting media (Vector Labs, Burlingame, CA) to prevent photobleaching.

### Imaging and data analysis

Control (contralateral, uninjured) retinas from both saline-treated (n=6) and HDAP2-treated (n=8) animals were analyzed as wholemounts to verify that HDAP2 does not affect RGC survival in the absence of injury, and to establish baseline regional RBPMS/ChAT ratios. Wholemounts were imaged using a Zeiss LSM 900 confocal microscope with a 10× objective by acquiring ~160 images (10% overlap) that were automatically stitched together using Airyscan processing. RGC counts of entire retinas were obtained using RGCode (Masin et al., 2021), to produce unbiased counts of RGCs.

RGC survival following ONC was assessed in 21 retinas (8 unoperated controls, 6 ONC untreated, 7 ONC + HDAP2 treated) with systematic sampling at 12 sites per retina for most animals (0.5, 1.5, and 2.5 mm from the optic nerve head in medial, temporal, dorsal, and ventral quadrants), yielding a total of 231 measurement sites. Retinal cells were imaged using an Olympus Fluoview 300 confocal microscope with an Olympus 40× oil immersion objective (NA 1.0). Laser intensity for the green channel was set at ~530V to enable comparison of RBPMS expression between treatment conditions.

In central regions, Z-stacks (1 μm steps) were collected through the ganglion cell layer to ensure imaging of RGCs above and below the middle focal plane, and through-focus projections were generated for cell counting. In midperipheral and peripheral retinal areas, single images were collected because ganglion cell layer cells were visible in a single focal plane. Each site was imaged separately using three wavelengths (488, 594, and 647 nm) and then merged in Photoshop to observe RGCs, starburst cells, and cell nuclei in the same fields.

Labeled cells in each image field (350 μm × 350 μm) were counted manually using ImageJ (NIH) while blinded to experimental condition and used to compute RGC densities and RBPMS:ChAT cell ratios for each location and condition. RBPMS expression in tissue was quantified using the “Analyze” feature in ImageJ to compute mean pixel intensity (in arbitrary fluorescence units (AU)) from confocal images.

### Optic Nerve Embedding and Transmission Electron Microscopy

Optic nerves were post-fixed in 1% osmium tetroxide for 30 minutes, dehydrated through a graded ethanol series (30%, 50%, 70%, 95%, 100%), and embedded in a 50-50 mixture of Spurr’s and EPON resins. Specimens were infiltrated with resin on a rotator at room temperature for 24 hours. Polymerization was performed in a 60°C oven for 18 hours. Ultrathin cross-sections were cut on a Leica Ultracut ultramicrotome at a thickness of 60 nm, mounted on 200-mesh copper grids, stained with uranyl acetate and lead citrate, and imaged at 80 kV using a Hitachi HT7800 transmission electron microscope. For each nerve (n=3 for each condition), 10 regions were imaged at 10,000 × magnification to count axons. Mitochondria within these fields were subsequently imaged at 50,000× magnification for detailed morphological assessment. Two investigators, blinded to the experimental condition, independently counted mitochondria to quantify mitochondrial density and morphology. Mitochondria were assessed using a 4-point quality grading scale: 1= inner and outer membranes were visible; cristae occupied 75 to 100% of the mitochondrial matrix; 2 = inner and outer membranes were visible; cristae occupied 50-74% of the mitochondrial matrix; 3 = membranes intact, noticeable reduction in cristae density 25-49 %, 4 = disrupted membranes, few or absent cristae.

### Statistical analysis

Retinal ganglion cell survival was evaluated using both absolute RGC densities and RBPMS/ChAT ratios. Total RGC counts from control wholemounts were compared between saline-treated and HDAP2-treated groups using Welch’s t-test and effect size was quantified using Cohen’s d.

Regional RBPMS/ChAT ratios across central, midperipheral, and peripheral retinal thirds from control animals (n=14) were analyzed using Friedman tests (non-parametric repeated measures) to assess for changes in ratios with retinal eccentricity.

For ONC experiments, statistical analyses employed mixed-effects linear models to account for repeated measurements within animals. Models were fitted using restricted maximum likelihood (REML) with animal ID as a random effect. Treatment group (control, ONC, ONC+HDAP2) and retinal region (center, midperiphery, periphery) were included as fixed factors, along with their interaction term. Post-hoc pairwise comparisons were performed using z-tests on model coefficients to compare treatment effects, with p-values adjusted using the Holm method for multiple comparisons.

ChAT+ amacrine cell densities were analyzed using the same mixed-effects modeling approach to verify stability of the internal reference population across treatment conditions

RBPMS fluorescence intensity between ONC + vehicle and ONC + HDAP2 groups was compared using t-tests, as these measurements represented single values per retinal region rather than repeated measures.

RGC densities in tables were calculated from mean RGC counts in 350 × 350 μm fields and converted to densities (cells/mm^2^) by multiplying by 1/(0.35 × 0.35) = 8.16. This allowed direct comparison with published data on RGC densities in mouse retinas.

Mitochondrial density and morphology grades were independently assessed by two investigators who were blinded to experimental condition. Inter-rater reliability was excellent (intraclass correlation coefficient ICC = 0.730, Pearson r = 0.895), and consensus values (average of both investigators) were used for all analyses. For each image, mitochondrial density was calculated as the number of mitochondria divided by the number of axons. Image-level densities were then averaged within each animal to obtain animal-level means (n=3 animals per group, 10 images per animal). Statistical comparisons among groups were performed using Kruskal-Wallis test followed by Mann-Whitney U tests for pairwise comparisons. Effect sizes were calculated as Cohen’s d based on animal-level means.

Data throughout are presented as mean ± SEM. Post-hoc power calculations confirmed adequate statistical power (1-β > 0.8) for detecting treatment effects on RBPMS/ChAT ratios at each retinal region. All statistical analyses were performed using the statsmodels package (v0.14.0) in Python v3.12, with significance defined as p < 0.05.

## Results

### HDAP2 is well tolerated at therapeutic concentrations

Toxicity testing showed a clear dose-dependent relationship between HDAP2 exposure and survival (Figure 1A). At the highest dose (300mg/kg, not shown), animals developed fulminant seizures and died within minutes. Doses of 100 mg/kg were also acutely lethal, and produced rapid sedation, seizure and death within 30 minutes. Intermediate doses of 50 mg/kg were better tolerated intitially, but all animals in this group died by day 9 of the 14-day treatment protocol. In contrast, mice treated with lower doses (3 to 10 mg/kg), tolerated daily injections for the full 14-day period with 100% survival. Animals in these lower dose groups maintained normal body weight and exhibited no behavioral abnormalities, indicating good overall tolerance to repeated systemic administration.

**Figure 1.**
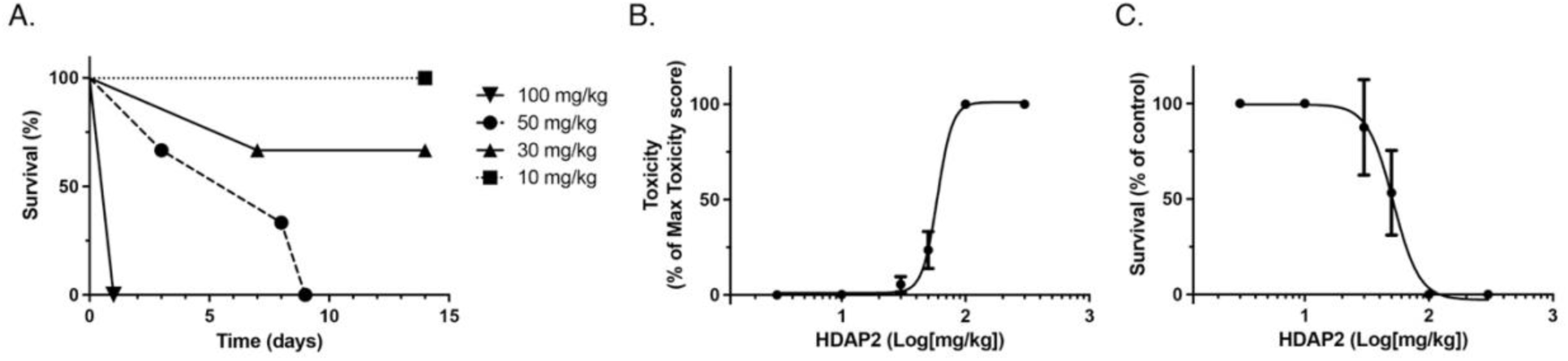
HDAP2 toxicity assessment and dose-response relationships. A. Kaplan–Meier survival curves of fixed-dose groups for intraperitoneal administration of HDAP2 (10–100 mg/kg/day for 14 days; n = 4–8 per group). Doses ≥100 mg/kg were acutely lethal, while doses below 30 mg/kg were tolerated with 100% survival. B. Dose–response curve for toxicity score indicates an LD_50_ of 59 mg/kg. Animals receiving <30 mg/kg exhibited no adverse side effects such as death, seizure, sedation or alterations in behavior (i.e. grooming, exploration) and body weight. C. Dose–response curve for survival of fixed-dose groups for the 14-day treatment protocol indicates an LD_50_ of 52 mg /kg. Data are presented as mean ± SEM for all assessments; P < 0.05 versus 3 mg/kg group (log-rank test for survival analysis).

Quantitative assessment of toxicity scores yielded an LD₅₀ of 55 mg/kg (95% CI: 46-59 mg/kg; Figure 1B-C). Kaplan-Meier survival curves produced a similar estimate of ~59 mg/kg. Based on these findings, the maximal tolerated dose for HDAP2 was established at 30 mg/kg. All ONC experiments used 3 mg/kg/day, well below the established toxic range.

### Distribution of HDAP2 peptide in the retina

To determine whether HDAP2 reaches retina neurons after systemic administration, we examined vertical sections and wholemounts from saline- and HDAP2-injected animals. Endogenous biotin produced faint streptavidin labeling in saline controls, with no detectable signal in RBPMS-positive RGCs (Figure 2A). In contrast, animals injected with 50 mg/kg HDAP2 showed strong streptavidin labeling throughout the ganglion cell layer, IPL, OPL, and photoreceptor inner segments (not shown) (Figure 2B). Labeling within RBPMS-positive cells displayed a perinuclear pattern with exclusion from nuclei, consistent with mitochondrial localization.

**Figure 2.**
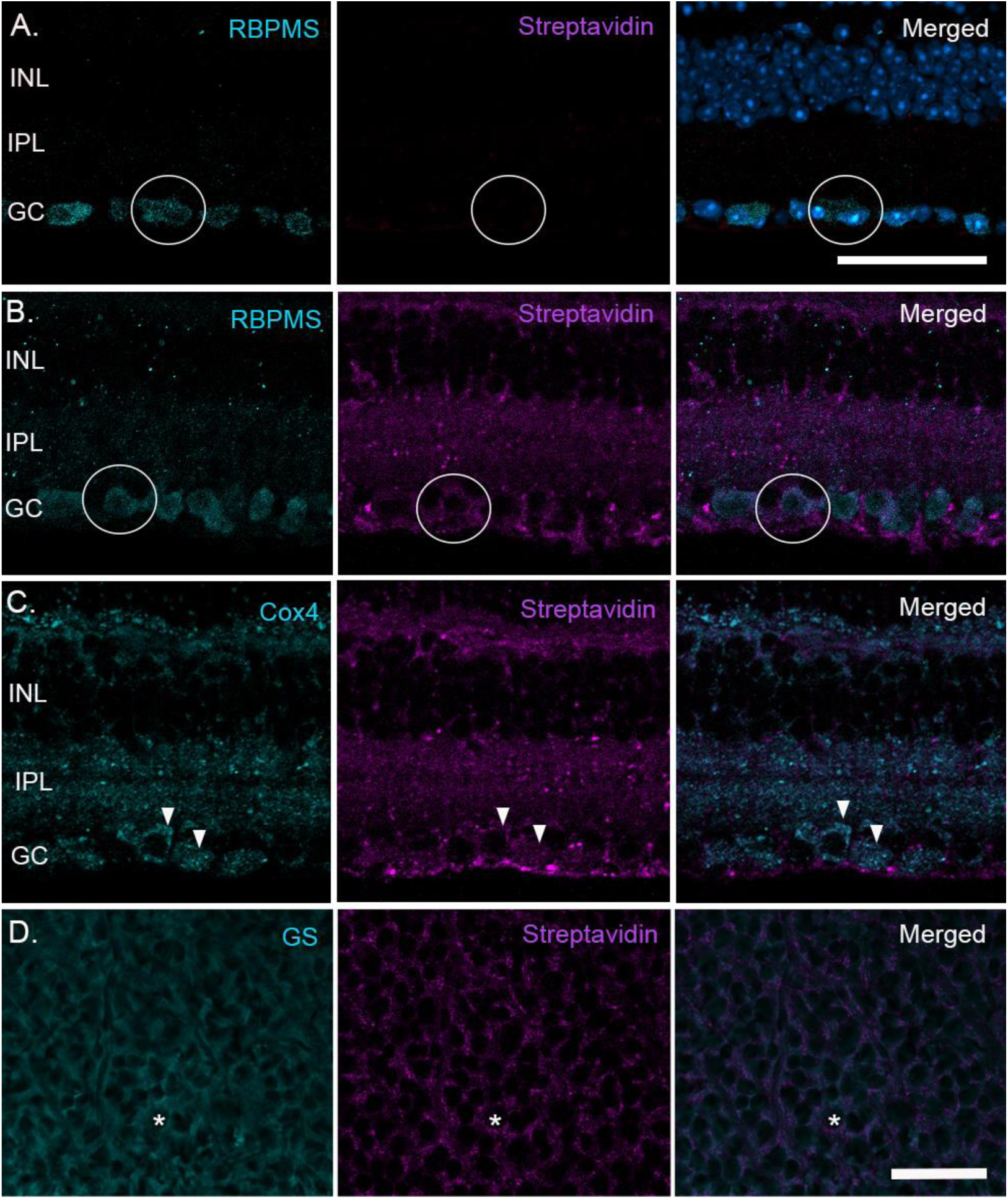
HDAP2 distribution and colocalization with RGCs, mitochondria, and Müller cells. Representative retinal images showing the distribution of endogenous biotin (A, middle panel) and biotinylated HDAP2 (B-D, middle panels) identified with Alexa Fluor 594-congugated streptavidin (magenta) in saline-injected (A) and HDAP2-injected Control retinas (B-D). Sections and wholemounts were double- and triple-labeled with RBPMS (cyan, to label RGCs), Cox4 (cyan, to label mitochondria), and ToPro3 (blue, to label nuclei) to assess colocalization. A. Minimal endogenous biotin labeling was observed in saline control tissue that did not overlap with RBPMS-labeled RGC somas (circles). B. Streptavidin labeling in HDAP2-treated retinas was robust throughout all retinal layers as well as in RGCs, identified by the perinuclear pattern in RBPMS-positive ganglion cells (circles). Labeling was characteristically excluded from nuclear regions (visible as unlabeled, black holes), which is consistent with mitochondrial localization. HDAP2 labeling is also present outside RGC somas in the GC fiber layer, indicating likely presence in Müller cells. C. Adjacent section co-stained with Cox4 (cyan, left) and streptavidin-Alexa Fluo594 (magenta, middle). The merged image (right) demonstrates extensive colocalization throughout retinal layers, including RGCs (arrowheads). D. Wholemounted retina labeled with glutamine synthetase (to label Müller cells) and streptavidin (middle) to show HDAP2 colocalization with Müller cells (one example marked with asterisks). Retinal layers are labeled: GC, ganglion cell layer; INL, inner nuclear layer; IPL, inner plexiform layer. Scale bars: 50 μm.

To confirm mitochondrial targeting, adjacent sections were co-labeled with Cox4 and streptavidin. The two signals showed extensive overlap across retinal layers and within RGC somas (Figure 2C). Additional labeling of wholemounts with glutamine synthetase demonstrated that HDAP2 also localized to Müller cells (Figure 2D). Together, these experiments show that HDAP2 crosses the blood-retina barrier and distributes broadly to retinal neurons and glia, including localization consistent with mitochondria-rich regions.

### HDAP2 does not alter RGC number or retinal organization in uninjured retinas

To assess potential off-target toxicity in uninjured tissue, we compared control retinas from animals treated with saline (n=6) or HDAP2 (n=8). RBPMS labeling showed normal RGC morphology and distribution in both groups (Figure 3A, top). Total RGC counts were comparable (49,417 ± 4,412 cells; p = 0.757, Cohen’s d = 0.171; Figure 3B), and both groups displayed the expected center-to-periphery density gradient (3,883 ± 60 to 2,295 ± 126 cells/mm^2^).

**Figure 3.**
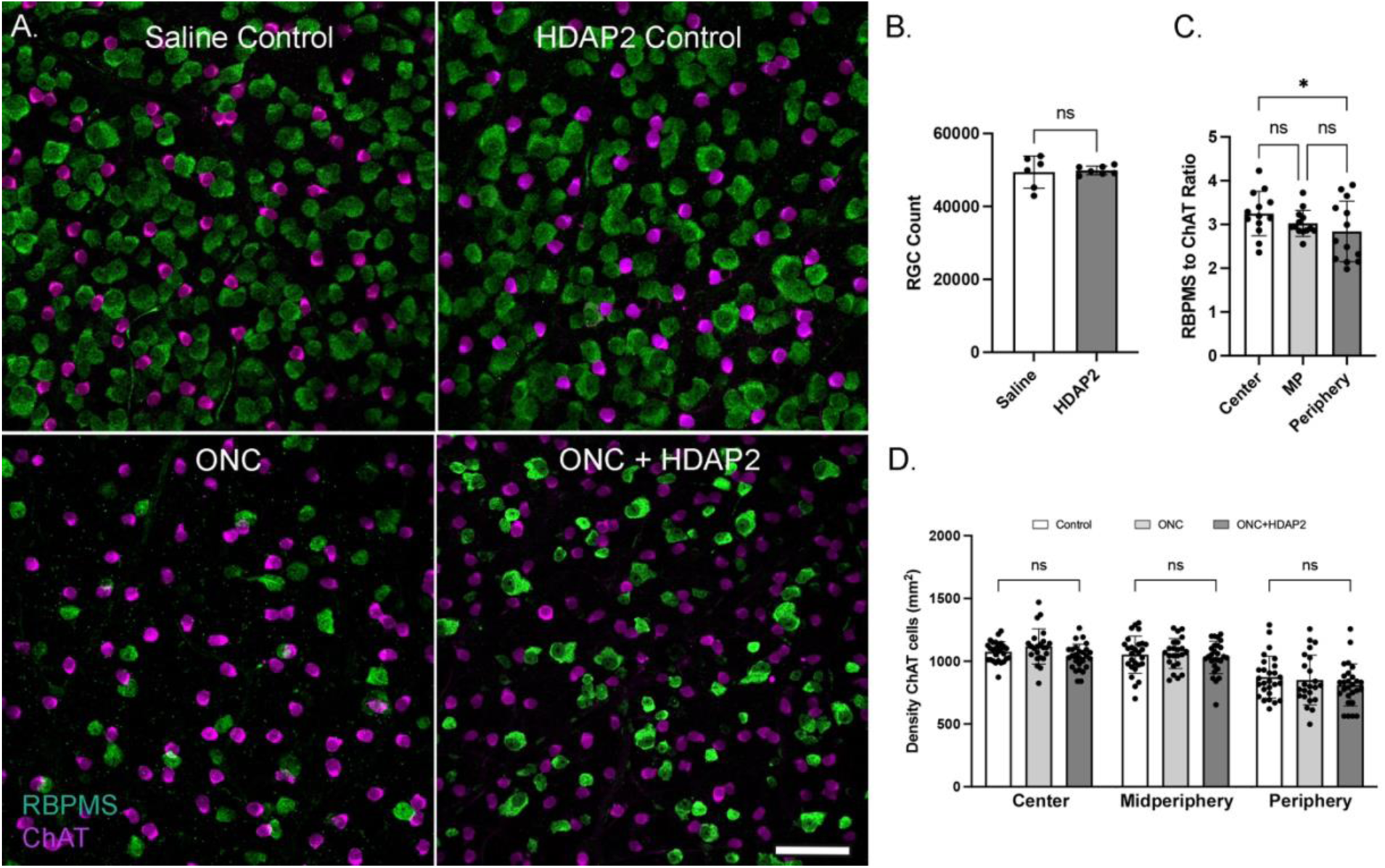
HDAP2 does not affect RGC counts in control retinas. A. Representative confocal images from peripheral retina showing RBPMS-labeled retinal ganglion cells (green) and ChAT-positive starburst amacrine cells (magenta) in control retinas (upper panels) and 14 days post–ONC (lower panels). In control retinas treated with vehicle control saline or HDAP2, RBPMS-labeled RGCs were indistinguishable in morphology and distribution to demonstrate that HDAP2 alone has no impact on normal retinal morphology or cell density. After ONC, many RGCs die in both HDAP2 and untreated conditions, but the patterns of ChAT labeling in the ganglion cell layer were unchanged. Scale = 50 µm. B. Total RGC counts (RBPMS-positive cells) in control retinas from untreated (n=6) and HDAP2-exposed (n=8) animals. No significant (ns) difference was observed between the groups (t-test, p = 0.757). C. Regional RBPMS/ChAT ratios across retinal regions for control wholemounts (n=14). Box plots show significant regional variation (Friedman test, χ^2^ = 6.14, p = 0.046), with ratios decreasing from inner to outer regions. Individual data points represent individual animals. This gradient necessitates region-matched comparisons when analyzing RGC survival after injury. *p < 0.05, ns = not significant. D. ChAT-positive displaced amacrine cell densities across retinal regions, separated by treatment group (mean ± SEM). ChAT labeling shows the expected center-to-periphery density gradient (p < 0.0001, mixed-effects linear model) but remains stable across treatment conditions within each region (all treatment vs control comparisons, ns). ChAT-positive cells serve as a reliable internal reference population unaffected by ONC or HDAP2 treatment. Control (n=8 animals), ONC (n=6 animals), ONC+HDAP2 (n=7 animals). ns = not significant.

Because RGC densities vary regionally, we also examined RBPMS/ChAT ratios in control wholemounts to establish the baseline distribution. Starburst amacrine cells, which survive ONC (Berry et al., 2015; Jakobs et al., 2005), display relatively uniform ChAT labeling across retinal regions (Jeon et al., 1998). Their ratios with RGCs showed modest but significant regional differences (Friedman test, p = 0.046), decreasing toward the periphery (3.62 ± 0.06 to 2.67 ± 0.14; Figure 3C). Overall, ChAT+ displaced amacrine cells, were stable across treatment groups and retained their normal regional gradient (Figure 3D). These results confirm that HDAP2 does not affect RGC survival or starburst amacrine cell density in the absence of injury and supported the use of region-matched analyses for ONC experiments.

### HDAP2 attenuates RGC loss following optic nerve crush

As expected, ONC caused severe RGC degeneration, with untreated animals losing 86.3% of RGCs by 14 days post-injury (Table 2A, Figure 4A). Average RGC densities decreased from approximately 3,269 cells/mm^2^ in control retinas to just 446 cells/mm^2^ in untreated ONC retinas. In striking contrast, HDAP2-treated animals maintained significantly higher RGC survival, with densities of 629 cells/mm^2^, representing a 40.8% improvement over untreated ONC animals (Table 2A, Figure 4A).

**Figure 4.**
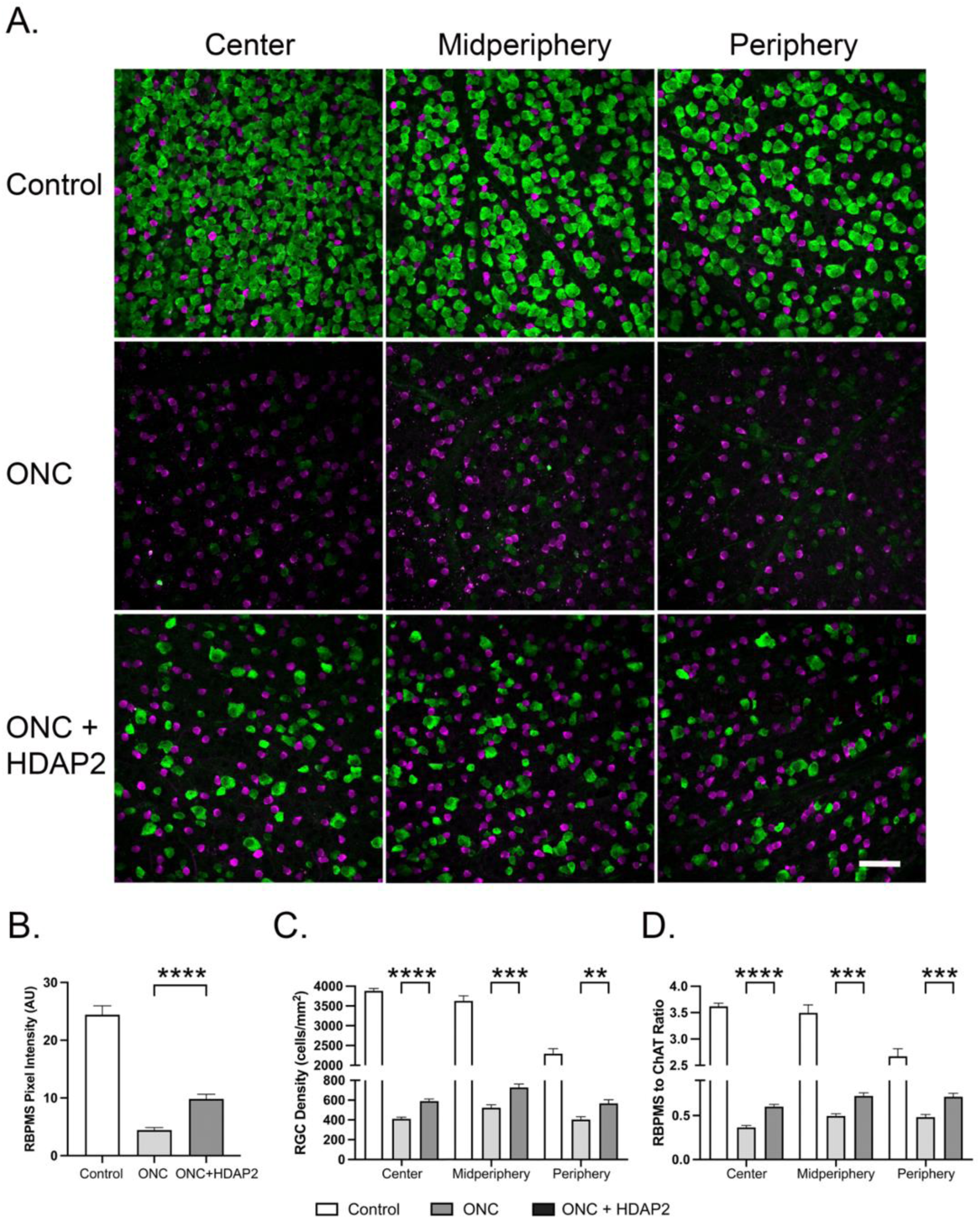
HDAP2 treatment enhances RGC survival following optic nerve crush. A. Retinal wholemounts labeled with RBPMS (green) to identify RGCs and choline acetyltransferase to label starburst amacrine cells (ChAT, magenta) from control, ONC, and ONC + HDAP2 treatment groups at 14 days post-injury. ONC induced extensive loss of RGCs compared to control retinas. However, HDAP2-treated animals (3 mg/kg/day, IP for 14 days) showed 40% greater RGC density compared to untreated ONC controls. Note the preservation of larger cell bodies in HDAP2-treated retinas, suggesting preferential survival of alpha-type RGCs. Scale bar = 50 µm. Images are representative of n = 6–8 animals per group. All images were acquired using identical confocal settings (laser power, PMT gain, offset) to enable direct intensity comparison. The markedly brighter RBPMS signal in HDAP2-treated RGCs indicates elevated RBPMS expression, suggesting better-preserved cellular health compared to the dimmer signal in vehicle-treated RGCs. B. RBPMS expression intensity, quantified from confocal images using identical acquisition parameters, was more than doubled in HDAP2-treated ONC eyes relative to untreated ONC eyes (9.8 ± 0.78 vs. 4.4 ± 0.45 arbitrary units; p < 0.0001), demonstrating that HDAP2 preserves not only RGC survival but also cellular health in surviving neurons. C. Analysis of absolute RGC density across retinal eccentricities showed that HDAP2 significantly increased RGC survival across all retinal regions: +43.7% (central, p<0.0001), +38.8% (midperipheral, p=0.0005), and +40.4% (peripheral, p=0.003). D. Analysis of RBPMS:ChAT ratios across retinal eccentricities showed statistically significant increases with HDAP2 treatment in all regions. The increases were +52.8% (central, p<0.0001), +43.5% (midperipheral, p<0.001), and +46.0% (peripheral, p<0.001), indicating preferential preservation of RGCs. Error bars represent SEM. Data are from 6–8 animals per group, with most retinas contributing 12 measurement sites. ****p < 0.0001, ***p < 0.001, **p < 0.01, ns = not significant.

**Table 2:** RGC densities and ratios of RBPMS:ChAT labeling to assess neuroprotective effects of HDAP2 at 14 days following optic nerve crush (ONC). Table 2A reports absolute RGC densities (cells/mm^2^) across eccentricities.; Densities were calculated by converting mean RGC counts per 350 µm × 350 µm sampling region to mm^2^ units. HDAP2 treatment significantly increased RGC survival in all retinal regions. Data are presented as mean ± SEM.; *Percentage improvement was calculated as: ((ONC+HDAP2) − ONC)/ONC × 100. **P-values reflect post-hoc pairwise comparisons (ONC vs ONC+HDAP2) from mixed-effects models using t-tests on model-estimated means. Statistical significance was set at p<0.05. Table 2B presents RBPMS:ChAT ratios across retinal eccentricities (central, midperipheral, peripheral) in unoperated controls and in ONC retinas with and without HDAP2 treatment (3 mg/kg/day, IP for 14 days). Ratios were derived from mixed-effects model estimated means, with standard errors accounting for within-animal correlation. HDAP2 treatment significantly improved the ratio in all regions, indicating selective preservation of RGCs relative to the densities of ChAT-positive amacrine cells.

| Region | Control | ONC | ONC + HDAP2 | % Improvement* | P-value ** |
| --- | --- | --- | --- | --- | --- |
| **Central** | $3883.0 \pm 60.0$ | $411 \pm 18.0$ | $591.0 \pm 21.0$ | 43.8% | $p < 0.0001$ |
| **Midperipheral** | $3629.3 \pm 130.2$ | $523.9 \pm 29.3$ | $727.7 \pm 35.8$ | 38.9% | $p = 0.0005$ |
| **Peripheral** | $2295.4 \pm 125.6$ | $404.3 \pm 28.9$ | $567.2 \pm 36.8$ | 40.3% | $p = 0.003$ |

B. RBPMS:ChAT ratios at different retinal eccentricities.
| Region | Control | ONC | ONC + HDAP2 | % Improvement* | P-value** |
| --- | --- | --- | --- | --- | --- |
| **Central** | $3.620 \pm 0.058$ | $0.375 \pm 0.021$ | $0.573 \pm 0.020$ | 52.8% | $p < 0.0001$ |
| **Midperipheral** | $3.496 \pm 0.149$ | $0.494 \pm 0.026$ | $0.709 \pm 0.034$ | 43.5% | $p < 0.001$ |
| **Peripheral** | $2.672 \pm 0.144$ | $0.483 \pm 0.029$ | $0.705 \pm 0.038$ | 46.0% | $p < 0.001$ |

Qualitative assessment of retinal wholemounts revealed distinct differences in RGC populations across treatment groups. In control retinas, RBPMS labeling was robust across all regions, displaying the expected center-to-periphery gradient with highest cell counts centrally and gradual decline toward the periphery (Figure 4A, control). Following ONC, there was severe loss of RBPMS signal and dramatic reduction in surviving RGCs (Figure 4A, ONC). HDAP2 treatment did not prevent this overall pattern of degeneration, but preserved substantially more RGCs compared to untreated retinas (Figure 4A, ONC+HDAP2).

Beyond simply preserving more cells, HDAP2 treatment appeared to maintain cellular health. Surviving RGCs in HDAP2-treated animals showed larger, healthier-appearing somas and more than double the RBPMS fluorescence intensity compared to untreated ONC retinas (9.8 ± 0.78 vs. 4.4 ± 0.45 AU, p < 0.0001), suggesting enhanced cellular health and function in HDAP2-protected cells (Figure 4B).

Quantitative analysis confirmed that HDAP2 provided consistent neuroprotection across all retinal regions. We assessed RGC survival using both absolute cell density measurements and RBPMS/ChAT ratios, which normalize RGC counts to the stable population of ChAT+ displaced starburst amacrine cells in each region (see Methods and Figure 3C, 3D). Both measures revealed the same pattern of robust protection. In the central retina, HDAP2-treated animals retained 590 ± 26 cells/mm^2^ compared to 410 ± 18 cells/mm^2^ in untreated ONC retinas, a 43.7% increase (p<0.0001), corresponding to an RBPMS/ChAT ratio increase from 0.375 ± 0.021 to 0.573 ± 0.020, a 52.8% improvement (p<0.0001; Tables 2A, 2B). Similarly, the midperipheral region showed 38.8% higher cell density (728 ± 36 vs. 524 ± 29 cells/mm^2^, p=0.0005) and 43.5% higher RBPMS/ChAT ratios (0.709 ± 0.034 vs. 0.494 ± 0.026, p<0.001). The peripheral retina demonstrated 40.4% increased density (567 ± 37 vs. 404 ± 29 cells/mm^2^, p=0.003) and 46.0% higher ratios (0.705 ± 0.038 vs. 0.483 ± 0.029, p<0.001). All regional comparisons were statistically significant (Figure 4C, 4D; Tables 2A, 2B), demonstrating that HDAP2 significantly attenuates RGC loss following optic nerve injury across the entire retina.

### HDAP2 reduces mitochondrial loss and axonal degeneration in the optic nerve

To determine whether HDAP2’s neuroprotective effects involved protection of axons and preservation of mitochondrial populations, we quantified mitochondrial density and morphology in optic nerve cross-sections using transmission electron microscopy (TEM). Untreated crushed nerves had a 29% decrease in axon number, with surviving axons showing extensive ultrastructural damage characteristic of axonal degeneration, including myelin unwinding, disorganization of neurofilaments and microtubules, and electron-dense axoplasm (Figure 5A, middle panel). Moreover, these axons exhibited severe loss of mitochondria, with mitochondrial density reduced by 87% compared to uninjured controls (0.025 ± 0.012 vs. 0.185 ± 0.017, mean ± SEM, n=3 animals per group; Kruskal-Wallis p = 0.039, Cohen d = 6.32; Figure 5C) and many images from untreated crushed nerves containing no visible mitochondria (37% of images vs. 0% in controls). When mitochondria were identified, they had recognizable morphology, even when the nerve itself appeared to show characteristics of neurodegeneration (Figure 5A, middle panel; Figure 5D).

**Figure 5.**
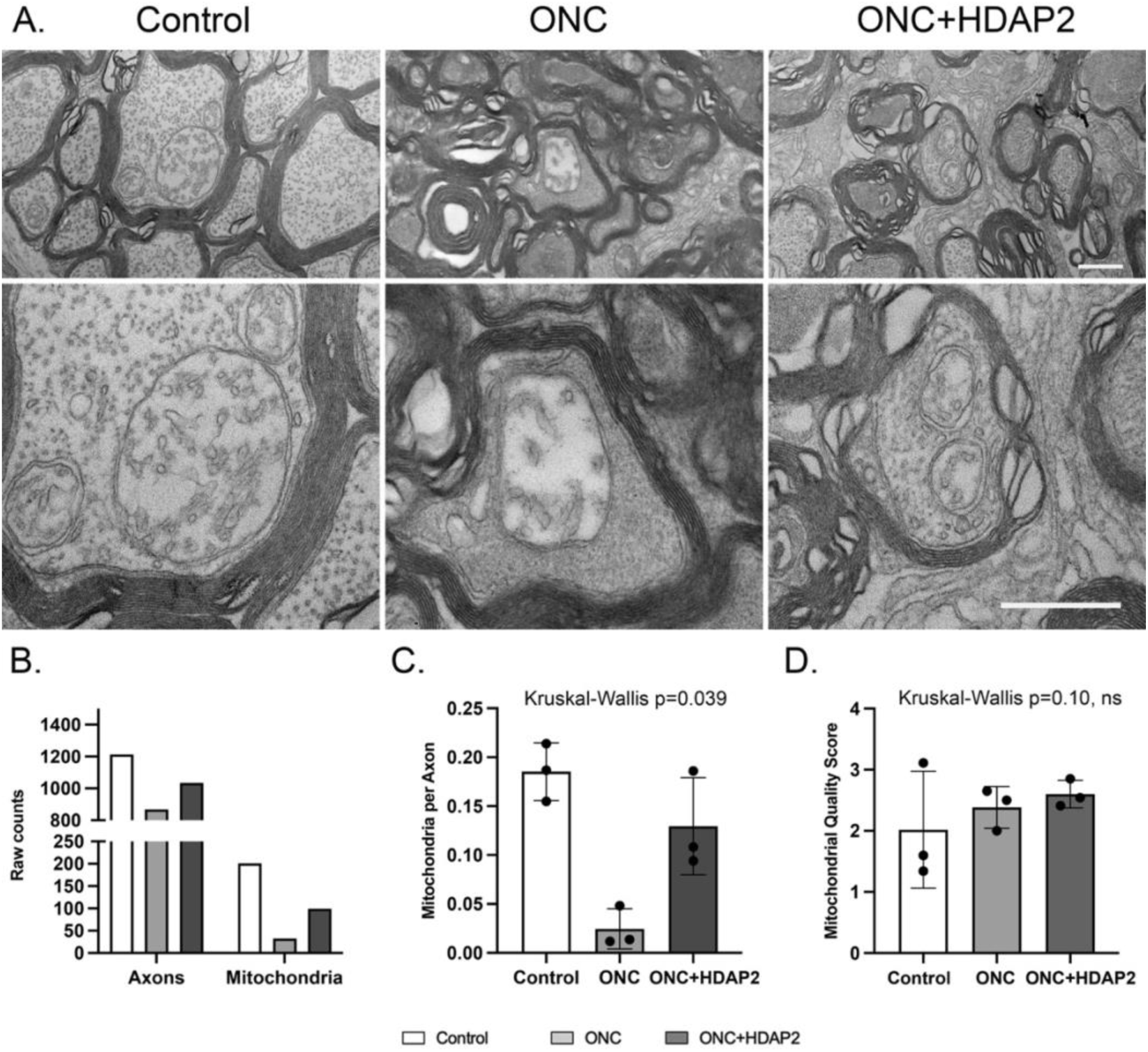
HDAP2 preserves axonal mitochondria following optic nerve crush. A. Representative transmission electron micrographs of optic nerve cross-sections 14 days post-injury. Upper panels: axonal cross-sections showing mitochondrial distribution (50,000×). Lower panels: individual mitochondria enlarged to demonstrate cristae preservation and membrane integrity. Scale bars = 500 nm B. Total axons and mitochondria quantified across all images. Data represent cumulative counts from 30 images per condition (10 images per animal, 3 animals per group). ONC reduced axon counts by 28% (867 vs 1213) and mitochondrial counts by 84% (32 vs 201) compared to control, while HDAP2 treatment provided substantial protection with counts approaching control levels. C. Mitochondrial density per axon by condition. Each data point represents the mean from one animal (average of 10 images). Bars show group mean ¬± SEM (n=3 animals per group). Groups differed significantly overall (Kruskal-Wallis p = 0.039). Effect sizes (Cohen’s d): Control vs Untreated = 6.32, Untreated vs HDAP2 = 2.77. HDAP2 treatment preserved mitochondrial density at 70% of control levels, representing 5.3-fold higher density than untreated ONC. D. Mitochondrial morphology grade (1=best preservation, 4=severe damage). Each point represents one animal. Bars show group mean ± SEM (n=3 animals per group). No significant difference in the quality score was noted among groups (Kruskal-Wallis p = 0.010).

In contrast, HDAP2-treated nerves showed partial but substantial axonal protection, with 19% more axons than untreated ONC (1034 vs 867 axons across all images, Figure 5B). While HDAP2-treated axons also exhibited signs of injury-related degeneration, healthy-appearing axons with preserved mitochondrial populations and relatively intact ultrastructure were consistently present throughout the nerve, unlike untreated ONC nerves where such preserved axons were rarely observed (Figure 5A, right panel). Mitochondrial density in HDAP2-treated crushed nerves was 0.129 ± 0.029, which represented a 5.3-fold higher density than vehicle-treated crush and was approximately 70% of uninjured control levels (Figure 5B), demonstrating a marked protective effect (Cohen d = 2.77 for HDAP2 vs. untreated ONC). While HDAP2 did not fully restore mitochondrial density to control levels (p = 0.20), it provided dramatic preservation compared to untreated injury. This mitochondrial preservation occurred even in axons showing other signs of injury, suggesting that HDAP2 specifically protects mitochondrial integrity within a degenerating axonal environment.

Morphological analysis of surviving mitochondria revealed similar ultrastructural characteristics across experimental groups. In general, control axons had mitochondria with the lowest degeneration scores, but the distribution of quality ratings did not differ significantly among conditions: 2.02 ± 0.55 (control), 2.36 ± 0.20 (ONC), and 2.60 ± 0.13 (ONC+HDAP2) (Figure 5D). These findings indicate that optic nerve crush injury drives mitochondrial loss rather than progressive degradation of surviving organelles, suggesting that damaged mitochondria are efficiently removed through quality control mechanisms such as mitophagy and that HDAP2 treatment preserves the population of functional, structurally intact mitochondria.

## Discussion

Mitochondrial dysfunction is a major driver of RGC loss following optic nerve injury. In this study, we found that the mitochondria-targeted peptide HDAP2 significantly attenuated RGC loss after optic nerve crush, improving survival by approximately 40% across all retinal regions. These findings were consistent across analysis methods, using both RGC densities and RBPMS:ChAT ratios, indicating robust treatment effects independent of regional cell density differences. HDAP2 also maintained substantially higher mitochondrial density in optic nerve axons after injury, demonstrating that its effects extend to the ultrastructural level.The therapeutic dose used here (3 mg/kg/day) was well below the maximal tolerated dose, which supports a wide safety margin for future translational development. Although the present experiments focused on the retina, the pathways involved, mitochondrial loss, axonal degeneration, and membrane destabilization, are shared across CNS projection neurons and may have broader relevance in traumatic axonal injury.

### RGC survival following ONC

We evaluated the survival of RGCs by comparing the absolute densities of RGCs in HDAP2 and untreated conditions, as well as by analyzing ratios of RBPMS-positive RGCs to displaced starburst amacrine cells labeled by ChAT. The latter metric was useful for comparing cell survival of small populations that are independent of retinal eccentricity. Since starburst amacrine cells lack axons, they survive ONC and can be used as a stable internal control. Our analysis confirmed that the densities of ChAT+ cells remained stable across all treatment groups and retinal regions, with no significant differences between control, ONC, and ONC+HDAP2 conditions (all comparisons p > 0.3). Baseline RBPMS:ChAT ratios did however decrease from center to peripheral regions (central: 3.62, midperipheral: 3.50, peripheral: 2.67; p = 0.046), which challenges the assumption that these ratios remain constant across retinal eccentricities (Jeon et al., 1998). This regional variation necessitated region-matched statistical comparisons to accurately assess treatment effects.

Following ONC, we observed that HDAP2 treatment significantly improved RBPMS/ChAT ratios in all retinal regions. In central retina, the ratio increased by 52.5% in HDAP2-treated animals compared to untreated ONC controls, with similar improvements in midperipheral (43.5%) and peripheral (46.0%) regions (all p < 0.0001; Table 2B). These findings closely align with absolute density measurements, which increased RGC survival by 40-44% across all regions, to provide strong evidence for HDAP2’s neuroprotective potential. The consistency between these two approaches strengthens confidence in our results and demonstrates that the observed protection reflects genuine RGC preservation rather than methodological artifacts.

### Regional Variation in RGC Vulnerability

The pattern of RGC loss following ONC was not uniform across retinal regions. Compared to control retinas, all regions experienced substantial cell death in ONC animals, but central retina showed proportionally greater loss (89% reduction from control) compared to peripheral regions (82% reduction). This differential vulnerability may reflect several factors. Central retina contains smaller RGC cell bodies with higher packing density (Dräger & Olsen, 1981; Jeon et al., 1998), which could potentially increase their metabolic demand per unit area. Under metabolic or ischemic stress, smaller axons with proportionally fewer mitochondria (Barron et al., 2004; Perge et al., 2009) may have limited energy reserves to sustain ATP generation, making them more vulnerable to energy failure and apoptotic signaling cascades (Kong et al., 2009). Additionally, central RGCs lie in closer proximity to the optic nerve head and crush site, where injury-generated inflammatory mediators and oxidative stress are most pronounced (Howell et al., 2011; Williams et al., 2013). Together, these factors may contribute to the greater vulnerability of central RGCs after ONC.

The patterns of regional protection may also depend in part on cells other than RGCs. Müller glia are critical providers of metabolic and trophic support to RGCs, such as with K⁺ buffering, glutamate–glutamine cycling, lactate shuttling, and antioxidant defense, and their function is highly reliant on mitochondrial integrity (Bringmann & Wiedemann, 2011). In contrast to RGCs, Müller cells are fairly evenly distributed throughout the retina (Wang et al., 2017). Consequently, the RGC to Müller cell ratio is higher in the central retina and lower in the peripheral retina, which may place greater metabolic demands on centrally located Müller cells, and may contribute to regional vulnerability. Since HDAP2 is expressed in Müller cells (Figure 2D), stabilization of mitochondrial in glia may indirectly lead to support of RGCs, particularly in areas of high neuronal activity. A direct evaluation of mitochondrial support in glial cells will be important for understanding their contributions to RGCs under conditions of cellular stress.

Despite regional differences in baseline vulnerability, HDAP2 provided remarkably consistent RGC attenuation across all retinal areas, with survival improvements ranging from 39-44%. This uniform protective effect suggests that HDAP2 acts through a fundamental mechanism common to all RGC populations rather than by targeting cell-specific vulnerabilities (Catalani et al., 2023; Muench et al., 2021). Given HDAP2’s mitochondrial-targeting properties, the most conservative explanation is that the peptide stabilizes mitochondrial function and preserves bioenergetic capacity in stressed RGCs throughout the retina. This interpretation is supported by previous studies demonstrating that mitochondria-targeted antioxidants and peptides can preserve ATP production, reduce reactive oxygen species generation, and prevent mitochondrial membrane permeabilization, which are key early events in RGC apoptosis following optic nerve injury (Birk et al., 2013; Iomdina et al., 2015; Wu et al., 2019).

### Maintenance of mitochondrial populations as a potential mechanism

The RGC preservation observed with HDAP2 treatment was accompanied by robust protection of axonal mitochondrial populations. Transmission electron microscopy revealed that ONC caused an 87% reduction in mitochondrial density, while HDAP2 treatment attenuated this loss, maintaining mitochondrial numbers at substantially higher levels. Despite injury, the morphological quality of surviving mitochondria was similar between untreated and treated groups, suggesting HDAP2 mainly reduces mitochondria elimination rather than alters the structural integrity of remaining organelles. This scenerio aligns with prior reports that ONC and related optic neuropathies trigger mitochondrial dysfunction and selective degradation of damaged mitochondria via mitophagy or autophagic mechanisms (Kang et al., 2019; Muench et al., 2021). The ability of HDAP2 to counteract mitochondrial loss is consistent with its design to stabilize mitochondrial membranes and mitigate injury-induced organelle elimination (Catalani et al., 2023; Liang et al., 2024). These findings directly link mitochondrial population preservation to RGC survival and are consistent with HDAP2’s mechanism of binding cardiolipin to stabilize mitochondrial membranes and prevent their elimination (Birk et al., 2025).

The maintenance of mitochondrial populations by HDAP2 likely contributes to improved neuronal survival through several complementary mechanisms. Mitochondria are essential for sustaining axonal ATP production, buffering intracellular calcium, and preventing oxidative stress–induced injury through maintenance of membrane potential (Nicholls & Budd, 2000; Schon & Przedborski, 2011). Disruption of these functions, including impaired ATP production, increased reactive oxygen species generation, and mitochondrial membrane permeabilization, represents a key early event in RGC apoptosis following optic nerve injury (Nicholls & Budd, 2000; Muench et al., 2021). Given HDAP2’s mitochondrial-targeting properties and its demonstrated preservation of mitochondrial populations, the peptide likely stabilizes these critical functions in stressed RGCs. This mechanism is consistent with other mitochondria-targeted therapeutic interventions that have shown neuroprotective efficacy in models of glaucoma and optic nerve injury (Nascimento-Dos-Santos et al., 2020; Wu et al., 2019; Tribble et al., 2021).

### Future Directions

Several key questions remain for future investigation. Dose-response studies should determine whether higher doses of HDAP2 (while remaining within the safety margin) could provide enhanced neuroprotection. Additionally, functional assessments, including electroretinography, optokinetic tracking, and visual evoked potentials, are needed to determine whether preserved RGCs retain function. The efficacy of earlier intervention (prior to injury) or longer treatment durations also warrants exploration to better understand the temporal window and sustained delivery requirements for HDAP2’s protective mechanism. Detailed mechanistic studies are needed to determine whether HDAP2 preserves mitochondrial populations by stabilizing existing organelles, promoting biogenesis, or inhibiting mitophagy.

## Abbreviations

ATP: adenosine triphosphate
ChAT: choline acetyltransferase
GCL: ganglion cell layer
HDAP2: high-density aromatic peptide
INL: inner nuclear layer
IP: intraperitoneal
IPL: inner plexiform layer
IS: inner segment
LD_50_: mean lethal dose
NAD^+^: nicotinamide adenine dinucleotide
OCT: optical cutting temperature
OLS: ordinary least squares
ONC: optic nerve crush
ONL: outer nuclear layer
OPL: outer plexiform layer
RBPMS: RNA-binding protein with multiple splicing
RGC: retinal ganglion cell
ROS: reactive oxygen species
SEM: standard error of measure
TEM: transmission electron microscopy

## Acknowledgments

We thank Ms. Karen Manifold for her expert care of the animals and Dr. Andrew Lesniewski for statistical consultation.

## Author Contributions

Margaret MacNeil: Conceptualization, Formal analysis, Funding acquisition, Investigation, Project administration, Supervision, Writing—original draft, Writing review & editing; Sara Arain: Investigation, Formal analysis; Widnie Mentor: Investigation, Formal analysis; Virginia Garcia-Marin: Writing review & editing; Alexander Birk: Conceptualization, Formal analysis, Funding acquisition, Investigation, Writing review & editing. All authors reviewed and approved the final manuscript.

## Declaration of Competing Interest

The Research Foundation, CUNY has received a patent for HDAP2 therapeutic applications, with MAM and AB listed as inventors. At present, the patent holders have no financial holdings, licensing agreements, or commercial relationships related to HDAP2. SA and VGM declare no competing interests.

## Funding

This work was supported by grants from the PSC-CUNY (642000-00-52, MAM), National Institutes of Health (R16GM149428, MAM), Department of Defense HBCU/MI (W911NF-24-1-0322, VGM) and the Social Profit Network (AB).

## Notes

### Competing Interest Statement

The authors have declared no competing interest.

### Summary of Updates

This revised version of our manuscript on the mitochondria-targeted peptide HDAP2 and optic nerve injury includes several substantial improvements that strengthen the mechanistic evidence and interpretation of our findings. Major Additions: Transmission Electron Microscopy Data: We added a comprehensive TEM analysis showing that HDAP2-treated optic nerves retain significantly higher densities of intact mitochondria compared to injured controls. This ultrastructural evidence directly supports HDAP2's mechanism of preserving mitochondrial populations after axonal injury and provides a critical mechanistic link between peptide treatment and improved RGC survival. Refined Interpretation: We revised our characterization of the neuroprotective effect to more accurately reflect its magnitude. While HDAP2 treatment produces statistically significant 41% improvement in RGC survival, we now emphasize this as attenuation of cell loss rather than complete protection. Importantly, we highlight that surviving RGCs in treated eyes show doubled RBPMS fluorescence intensity, suggesting improved cellular health beyond mere cell number preservation. The manuscript title and running title were updated to "attenuates RGC loss" rather than "protects" to reflect this more nuanced interpretation. Discussion Enhancements: We expanded the mechanistic context regarding how cardiolipin stabilization contributes to maintaining mitochondrial integrity after injury, and provided clearer discussion of study limitations and future directions. Authorship: Widnie Mentor was added as an author in recognition of her contributions to the TEM experiments and analyses. These revisions substantially strengthen the mechanistic foundation of our work and provide a more accurate representation of HDAP2's neuroprotective effects in this optic nerve injury model.

